# Engineering Highly Homogenous Tetravalent IgGs with Enhanced Sperm Agglutination Potency

**DOI:** 10.1101/2020.04.27.064865

**Authors:** Bhawana Shrestha, Alison Schaefer, Elizabeth C. Chavez, Alexander J. Kopp, Timothy M. Jacobs, Thomas R. Moench, Samuel K. Lai

**Author notes:** Corresponding Author Samuel K. Lai, Division of Pharmacoengineering and Molecular Pharmaceutics, University of North Carolina at Chapel Hill, Marsico Hall 4213, 125 Mason Farm Road, Chapel Hill, NC 27599.

## Abstract

Millions of women avoid using available contraceptives and risk unintended pregnancies every year, due to perceived and/or real side-effects associated with the use of exogenous hormones. Naturally occurring anti-sperm antibodies can prevent fertilization in immune infertile women by limiting sperm permeation through mucus, particularly multivalent antibodies such as sIgA that offers robust agglutination potencies. Unfortunately, sIgA remains challenging to produce in large quantities and easily aggregates. Here, we designed two tetravalent anti-sperm IgGs with a Fab domain previously isolated from an immune infertile woman. Both constructs possess at least 4-fold greater agglutination potency and induced much more rapid sperm agglutination than the parent IgG while exhibiting comparable production yields and identical thermostability as the parent IgG. These tetravalent IgGs offer promise for non-hormonal contraception and underscore the multimerization of IgG as a promising strategy to improve existing mAb therapeutics.

---

Globally, over 40% of all pregnancies are unintended, which creates an enormous burden on healthcare systems.[1] For instance, in the U.S., nearly half of all pregnancies are unintended, resulting in an annual burden in excess of $20 billion per year.[2,3] Despite the availability of cheap and effective contraceptive methods, many women are dissatisfied with available contraceptive methods, particularly due to real and/or perceived side-effects associated with the use of exogenous hormones, including increased risks of breast cancer, depression, prolonged menstrual cycle, nausea and migraines.[4,5] Many women are also restricted from using estrogen-based hormonal contraceptives due to medical contraindications.[6–8] These realities strongly underscore the need for convenient non-hormonal contraceptives. Unfortunately, few options are currently available.

An effective non-hormonal contraceptive mechanism already exists in nature: anti-sperm antibodies (ASAs) in infertile women, particularly polyvalent immunoglobulins such as IgA, can arrest highly motile sperm in mucus and prevent sperm from permeating through mucus and reaching the egg.[9,10] At high sperm concentrations, ASA can agglutinate sperm into clusters that are too large to permeate through mucus.[11] At lower sperm concentration, ASA can also immobilize individual spermatozoa in mucus via multiple low affinity Fc-mucin bonds between sperm-bound ASA and mucins.[12,13] Vaginal delivery of ASA that sustains pharmacologically active doses of ASA in the female reproductive tract can make possible both consistent contraception, and rapid reversibility. Indeed, vaginal delivery of sperm agglutinating ASA exhibited considerable contraceptive efficacy in a rabbit model, reducing embryo formation by 95% in the highly fertile rabbit model.[14]

Owing to its prevalence at mucosal surfaces and strong agglutination potency due to the diametrically-opposite orientation of the 4 Fab domains, sIgA represents an attractive format to develop ASA-based contraceptive.[15] Unfortunately, this topical passive immunocontraception approach has never been advanced in humans, in part because of manufacturing challenges and stability issues with sIgA and IgM. In contrast, IgGs are highly stable and easy to produce, making IgG the most common antibody format for development of biologics. We hypothesized we could overcome the instability and production challenges of sIgA as well as the limited agglutination potency of IgG by creating IgG molecules with 4 Fabs, identical to sIgA. This led us to engineer two tetravalent (i.e. 4 Fabs) sperm-binding IgGs: Fab-IgG and IgG-Fab, based on linking an additional Fab to either the N- or C-terminus respectively, of a parent IgG using a flexible glycine-serine linker (Figure 1a). We designed our engineered Fab-IgG and IgG-Fab molecules to target a surface glycoprotein antigen, CD52g, a glycoform of CD52 that is unique to the male reproductive tract and present in abundant quantities on the surface of all sperm, but absent in all other tissues and women.[16,17]

**Figure 1.**
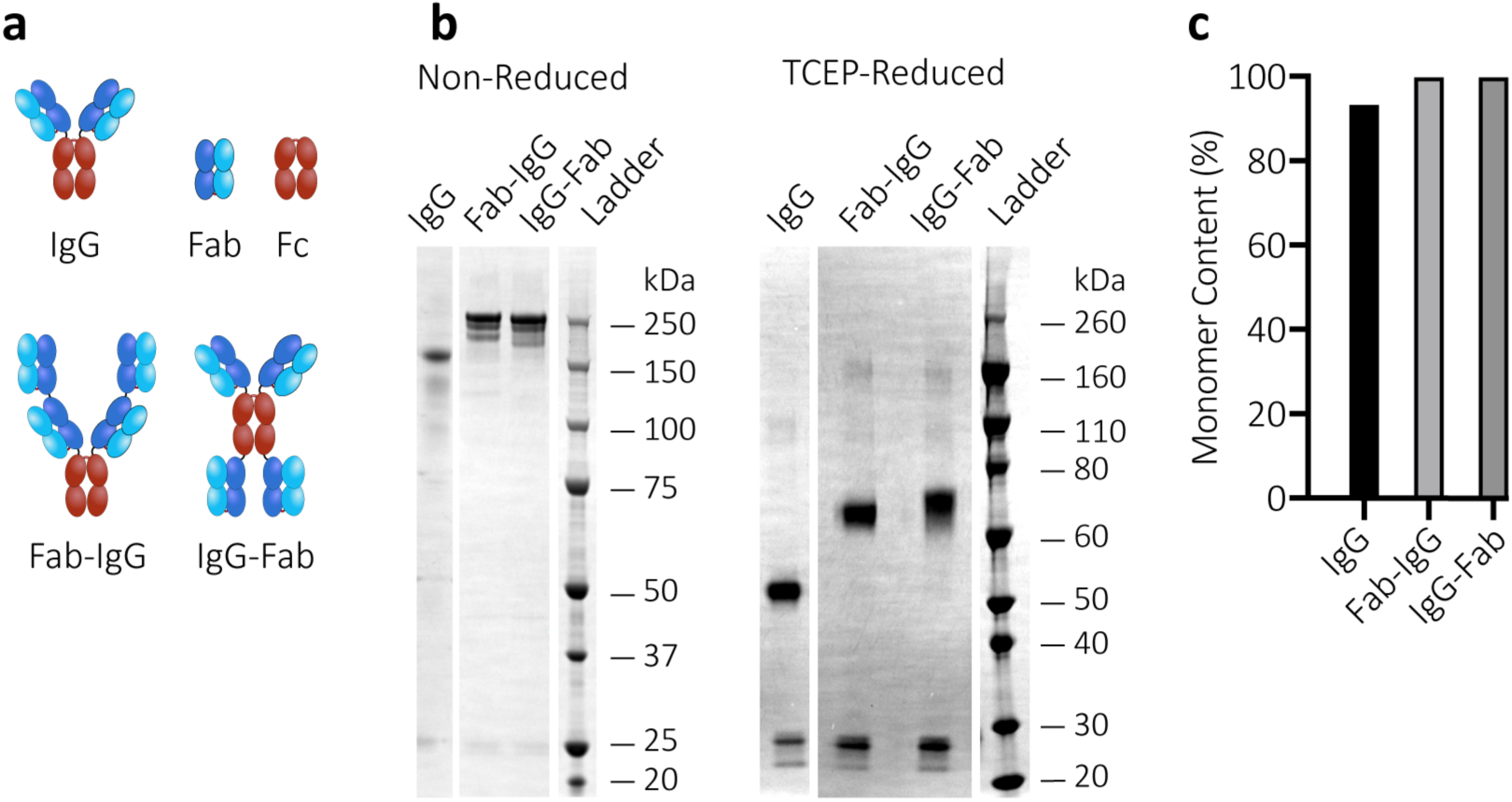
Production and characterization of tetravalent anti-sperm IgG antibodies. (a) Schematic diagrams of anti-sperm IgG, Fab-IgG and IgG-Fab. The additional Fab is linked to the N-terminal or C-terminal of parent IgG using flexible glycine-serine linkers. (b) Non-reducing and reducing SDS-Page analysis of the indicated Abs (1 μg) after expression in Expi293 cells and purification by protein A/G chromatography. Non-reducing SDS-Page showcases the total molecular weight of the Abs. Tris (2-carboxyethyl) phosphine hydrochloride (TCEP)-mediated reducing SDS-Page displays the molecular weight of individual heavy chain and light chain of Abs. The experiment was repeated independently two times with similar results. The SDS-PAGE image was adjusted for brightness and contrast using Fiji software. (c) Demonstration of the purity and homogeneity of the indicated Abs (50-100 μg) using Size Exclusion Chromatography with Multiple Angle Light Scattering (SEC-MALS) analysis. Y-axis indicates the total percentage of Abs representing their theoretical molecular weights.

Upon transient transfection in Expi293 cells, Fab-IgG and IgG-Fab expressed at comparable levels to the parent IgG (Figure S1a). The parent IgG, Fab-IgG and IgG-Fab all exhibited their expected molecular weights of 150 kDa, 250 kDa and 250 kDa, respectively (Figure 1b and Figure S1b). Surprisingly, while the parent IgG possessed a small fraction of aggregates that is commonly observed with IgGs, the tetravalent IgGs appeared completely homogeneous with no detectable aggregation (Figure 1c). We next evaluated the thermostability of the Fab-IgG and IgG-Fab using Differential Scanning Fluorimetry (DSF). Both constructs exhibited exceptional thermal stability, unfolding only at high temperatures of Tm_1_ (midpoint of unfolding of Fab and C_H_2) ≥ 71.1°C and Tm_2_ (midpoint of unfolding of C_H_3) ≥ 80°C, comparable to those of the parent IgG (Figure S1c). The incorporation of additional Fab appeared to have a stabilizing effect as Tm_1_ of Fab-IgG and IgG-Fab is slightly greater than Tm_1_ of native IgG. We next confirmed the binding of Fab-IgG and IgG-Fab to their sperm antigen using a whole sperm ELISA assay. Both constructs bound comparably to the human sperm as the parent IgG at 0.1 μg/mL (Figure S1d). We visually confirmed sperm agglutination by the parent IgG vs. tetravalent IgGs at their effective concentrations using scanning electron microscopy (Figure 2).

**Figure 2.**
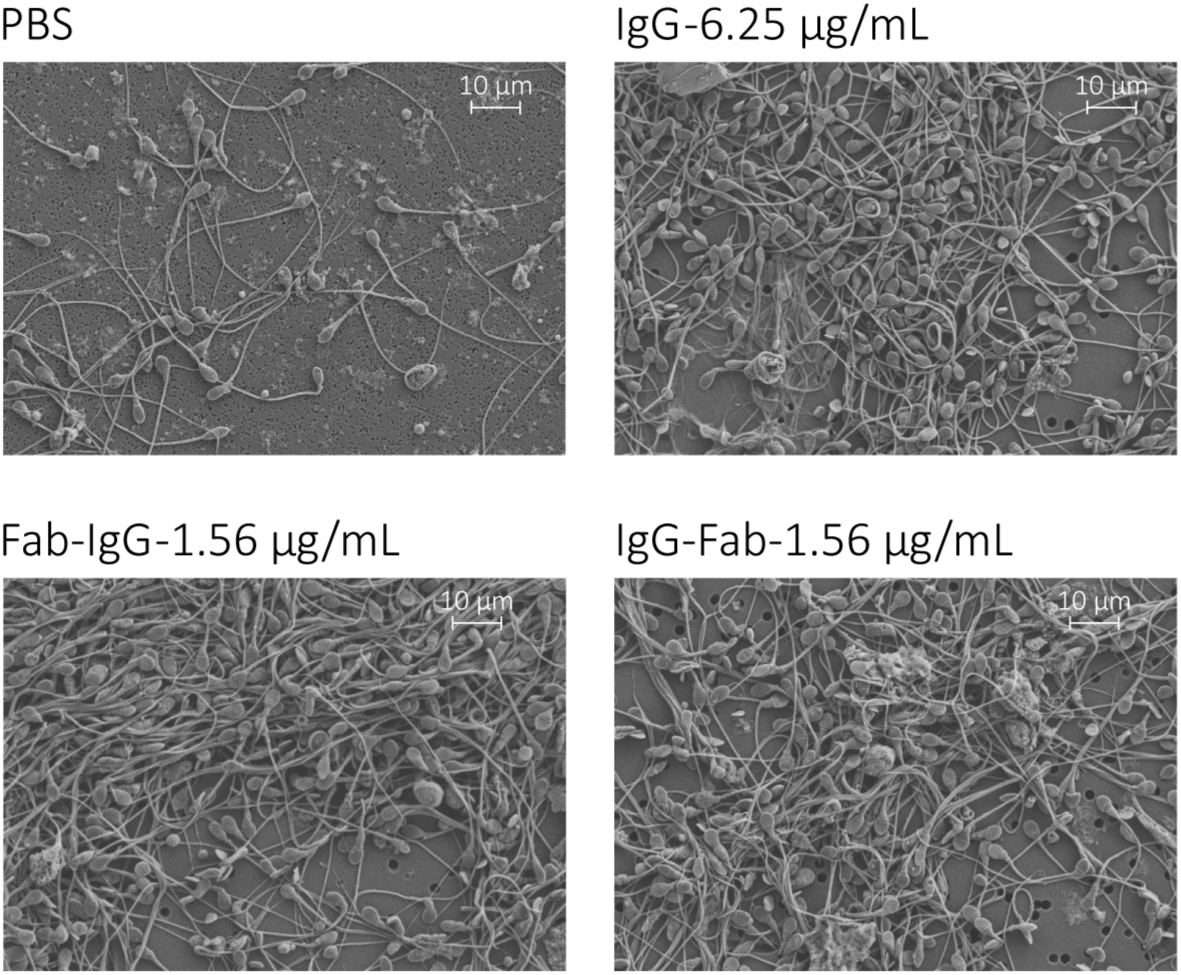
Scanning electron microscopy images of sperm agglutination. 20 million washed sperm were treated with IgG, Fab-IgG and IgG-Fab for 5 min and fixed with 4% PFA. Images were obtained at 2500X magnification. Scale bar, 10 μm.

Progressively motile (PM) sperm, due to their capacity to swim through mucus to reach the egg, is the key sperm fraction responsible for fertilization. We thus assessed the ability of Fab-IgG and IgG-Fab to agglutinate human sperm, using an in vitro sperm escape assay that quantified the number of PM sperm that escaped agglutination when treated with specific antibodies (Abs) vs the sperm washing media control determined by Computer Assister Sperm Analysis (CASA).[18,19] The assay was carried out at a concentration of 5 million PM sperm/mL, reflecting typical amounts of PM sperm in fertile males.[20,21] We found that Fab-IgG and IgG-Fab both exhibited markedly greater sperm agglutination potency than parent IgG. The minimal mAb concentrations needed to reduce PM sperm >98% was reduced from 6.25 μg/mL for the parent IgG to 1.56 μg/mL for both Fab-IgG and IgG-Fab (Figure 3a). Although both constructs failed to achieve >98% agglutination at 0.39 μg/mL, they were still able to reduce PM sperm by >80% (Figure 3b). In comparison, the parent IgG at 0.39 μg/mL offered no detectable reduction in PM sperm populations.

**Figure 3.**
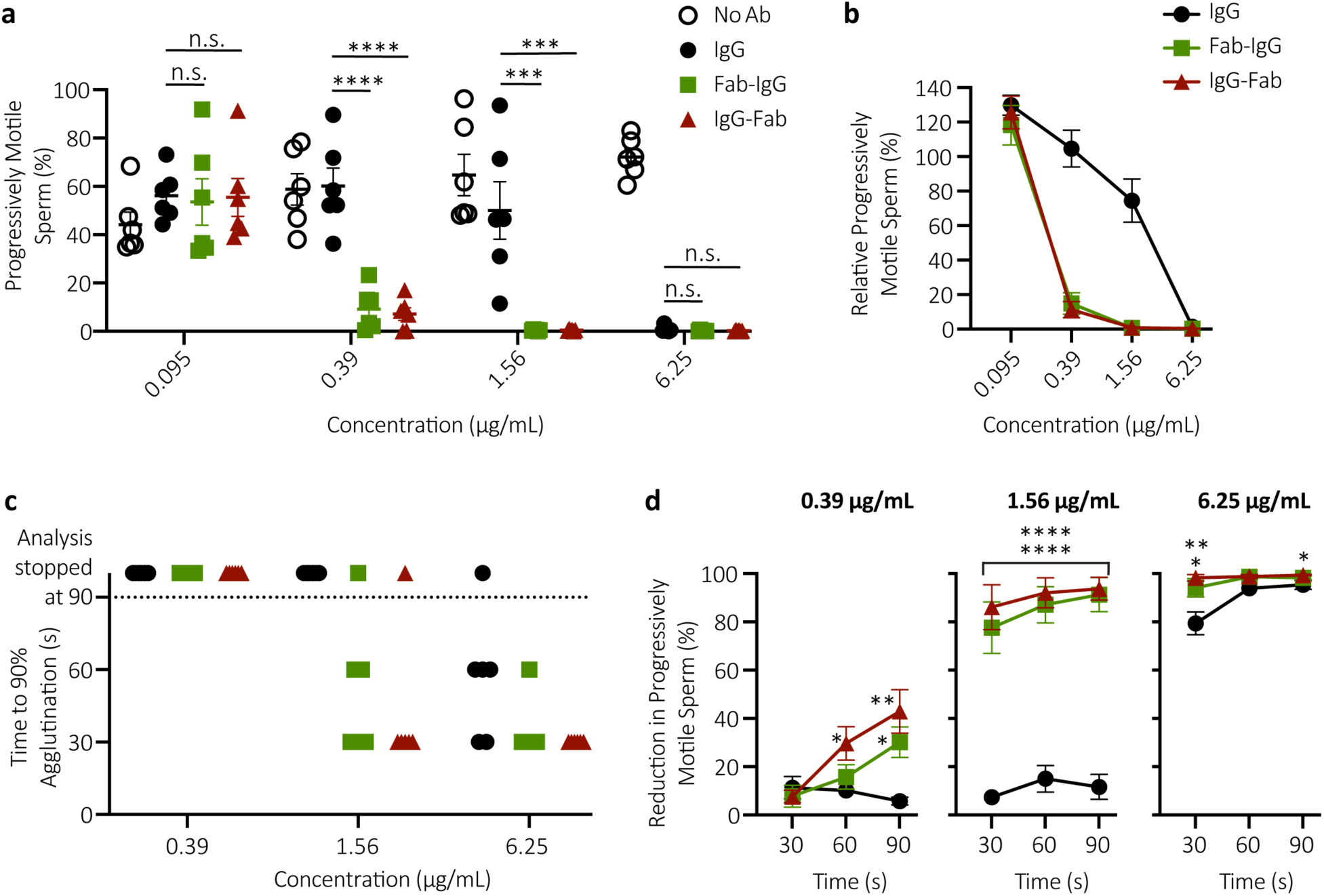
Multimerization markedly enhances the agglutination potency and kinetics of anti-sperm IgG antibodies. (a) Sperm agglutination potency of the parent IgG, Fab-IgG and IgG-Fab measured by the CASA-based quantification of the percentage of sperm that remains progressively motile (PM) after Ab-treatment compared to pre-treatment condition. (b) The sperm agglutination potency of the Abs normalized to the sperm washing media control. (c) Sperm agglutination kinetics of the parent IgG, Fab-IgG and IgG-Fab measured by the quantification of time required to achieve 90% agglutination of PM sperm compared to the sperm washing media control. (d) The rate of sperm agglutination determined by the reduction in percentage of PM sperm count at three timepoints after Ab-treatment compared to the sperm washing media control. Purified motile sperm at the final concentration of 5 × 10^6^ PM sperm/mL was used. Data were obtained from n = 6 independent experiments with at least n = 4 unique semen donors. Each experiment was performed in duplicates and averaged. P values were calculated using a one-way ANOVA with Dunnett’s multiple comparisons test. *P < 0.05, **P < 0.01, ***P < 0.001 and ****P < 0.0001. Lines indicate arithmetic mean values and standard error of mean.

For effective contraception, sperm must be stopped in mucus before they can swim through the cervix and access the uterus.[22] This suggests Abs that could agglutinate sperm more quickly should provide more effective contraception. Thus, we next quantified the sperm agglutination kinetics of both constructs using CASA by measuring the fraction of agglutinated and free PM sperm over time immediately after mixing washed sperm with different mAbs. At 6.25 μg/mL, the parent IgG reduced PM sperm by ≥90% within 90s in 5 of 6 semen samples; at 1.56 μg/mL, the parent failed to do so in all 6 of 6 samples (Figure 3c). In contrast, Fab-IgG and IgG-Fab achieved ≥90% agglutination within 30s in all 6 of 6 samples at 6.25 μg/mL, and within 60s in 5 of 6 samples at 1.56 μg/mL. Notably, the agglutination kinetics of Fab-IgG and IgG-Fab were markedly faster and more complete than the parent IgG at each Ab concentrations tested and across all timepoints (Figure 3d).

Earlier works have shown that IgA and IgG Abs can completely immobilize individual spermatozoa in the mucus by crosslinking antibody-bound spermatozoa to mucins; this is commonly referred to as the “shaking phenomenon’.[10] We have previously shown that multiple Fc present on Herpes-bound IgGs can form polyvalent adhesive interactions with cervicovaginal mucus (CVM), resulting in effective immobilization of the virus that in turn blocked vaginal Herpes transmission in mice.[12] Since the Fc region was conserved and identical in both tetravalent IgGs and parent IgG, we hypothesized that Fab-IgG and IgG-Fab constructs will trap individual spermatozoa in mucus similar to native IgG. We evaluated the muco-trapping potencies of the parent and tetravalent IgGs in CVM by multiple particle tracking, using a convolutional neural network to quantify the motion of fluorescently labeled sperm in mAb-treated CVM.[23] Both IgG-Fab and Fab-IgG reduced the fraction of PM sperm to a comparable extent as the parent IgG (Figure 4), indicating that Fc-mediated crosslinking remained unaffected by the addition of Fabs at both N- and C-terminus.

**Figure 4.**
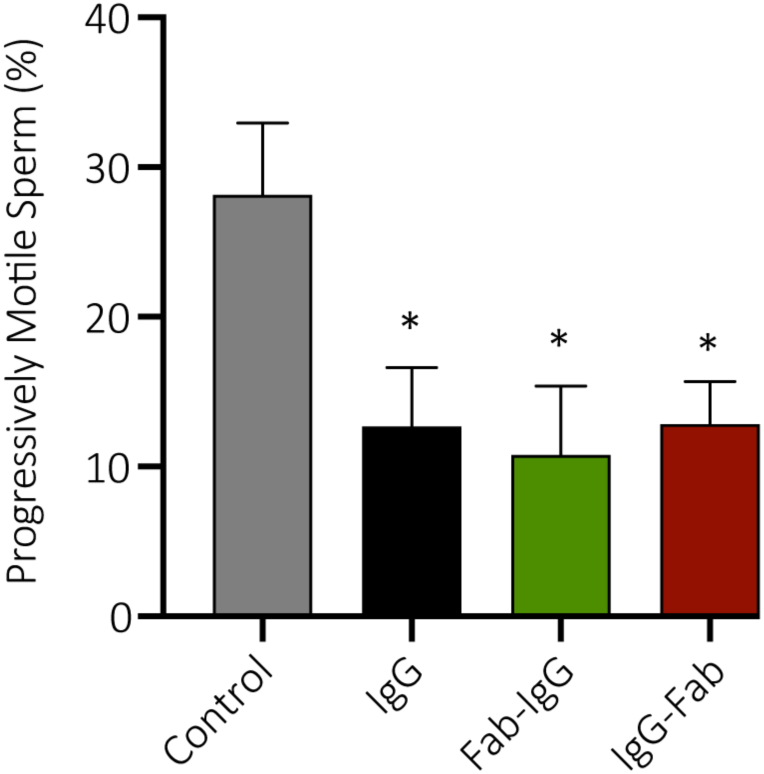
Tetravalent sperm-binding IgG constructs conserve the trapping potency of the parent IgG. The trapping potency of the indicated Abs measured by quantifying the percentage of fluorescently labeled PM sperm in Ab-treated CVM using neural network tracker analysis software. 25 μg/mL of Abs and purified motile sperm at the final concentration of 5.8 × 10^4^ PM sperm/mL were used. Data were obtained from n = 6 independent experiments with 6 unique combinations of semen and CVM specimens. P values were calculated using a one-tailed t-test. *P < 0.05. Lines indicate arithmetic mean values and standard error of mean.

Years ago, observations with naturally occurring immune infertility in women motivated the development of contraceptive vaccines.[24–27] Vaccines eliciting ASA offered considerable contraceptive efficacy, but the approach stalled due to unresolved variability in the intensity and duration of the vaccine responses in humans, as well as concerns that active vaccination might lead to permanent infertility.[28] In contrast, the local delivery of ASA to the vagina can overcome the key drawbacks of contraceptive vaccines by sustaining pharmacologically active doses of ASA and making possible both consistent contraception, and rapid reversibility. Unlike small molecule contraceptives, contraceptive mAbs should be exceptionally safe due to the specificity of targeting, particularly when binding to unique epitopes present only on sperm and not on expressed in female tissues. Safety is likely to be further enhanced by topical delivery: mAb delivered to mucosal surfaces such as the vagina are poorly absorbed into the systemic circulation,[29] and the vagina represents a poor immunization inductive site, with limited immune response even when vaccinating with the aid of highly immunostimulatory adjuvants.[30] Topical delivery also substantially reduces the overall mAb dose needed. Given the limited volume of secretions in the female reproductive tract (FRT), typically ≤1 mL in the vagina,[31] relatively high concentration of mAb locally can be achieved even with very limited total quantities of mAb dosed. In contrast, systemically dosed mAbs must contend with the large blood volume (∼5L), distribution to non-target tissues, natural catabolic degradation, and limited distribution into the FRT, including the vagina. The reduced quantities of mAb needed to sustain contraceptive levels in the FRT with vaginal delivery should translate to substantially lower amounts of total mAb needed, and consequently cost savings.

Many of the current multivalent Abs are bispecific or trispecific in nature and must content with potential mispairing of light and heavy chains. As a result, many such engineered Ab formats, such as single-chain variable fragment (scFv) or camel-derived nanobodies, involve substantial deviation from natural human Ab structure. scFv-based multivalent Ab constructs frequently suffer from low stability, heterogeneous expression, and decreased affinity and specificity stemming from the removal of the C_H_1/C_L_ interface present in a full-length Fab. The introduction of orthogonal mutations to facilitate heavy and light chain pairing can also substantially reduce mAb yield or overall stability. These limitations do not apply when generating monospecific multivalent IgGs, which can possess identical and full length human Fabs. Our strategy to covalently link additional Fabs to a parent heavy chain also contrasts from current multimerization strategies based on self-assembly of multiple IgGs based on Fc-mutations [32] or appending an IgM tail-piece, which often suffers poor homogeneity and stability.[33,34] The combination of fully intact human Fabs and covalent linkages likely contributes to the surprising thermal stability, homogeneity and bioprocessing ease of the multivalent IgGs developed here. We believe the IgG multimerization strategy presented here is likely a promising platform for developing mAbs where agglutination represents a critical effector function.

## Acknowledgements

This work was financially supported by the Eshelman Institute of Innovation (S.K.L.); The David and Lucile Packard Foundation (2013-39274; S.K.L); National Institutes of Health under grants R56HD095629 (S.K.L.), U54HD096957 (T.R.M. and S.K.L.), R43HD094454 (T.R.M.) and R44HD097063 (T.R.M.); National Science Foundation (DMR-1810168; S.K.L.); and PhRMA Foundation Graduate Fellowship (B.S.). Special thanks to Dr. Deborah O’Brien for providing the CASA instrument and assistance in setting up the CASA measurements. Special thanks to the UNC Macromolecular Interactions Facility for instruments used in SEC-MALS and DSF studies, and Microscopy Services Laboratory for instruments used in SEM experiments.

## Materials and methods

### Study design and ethics

All studies were performed following a protocol approved by the Institutional Review Board of the University of North Carolina at Chapel Hill (IRB-101817). Informed written consent was obtained from all male and female subjects before the collection of any material. Subjects were recruited from the Chapel Hill and Carrboro, NC area in response to mass student emails and posters.

### Cloning and production of the parent and tetravalent anti-sperm IgG antibodies

The variable heavy (V_H_) and variable light (V_L_) DNA sequences for anti-sperm IgG1 antibody (Ab) were obtained from the published sequence of H6-3C4 mAb.[11, 35] For light chain (LC) production, a gene fragment consisting of V_L_ and C_L_ sequences (Integrated DNA Technologies) was cloned into an empty mammalian expression vector (pAH, ThermoFisher Scientific). For parent IgG and Fab-IgG heavy chain (HC) production, V_H_ and V_H_/C_H_1-(G_4_S)_6_ linker-V_H_ gene fragments (GeneArt, ThermoFisher Scientific) were respectively cloned into mammalian IgG1 expression vector comprising of only C_H_1-C_H_2-C_H_3 DNA sequence. For IgG-Fab HC production, (G_4_S_)6_ linker-V_H_/C_H_1 was cloned into the parent IgG expression plasmid. The expression plasmids encoding HC and LC sequences were co-transfected into Expi293F cells using ExpiFectamine™ 293 Transfection reagents (Gibco). For IgG expression, HC and LC plasmid were co-transfected using a 1:1 ratio at 1 μg total DNA per 1 mL of culture. For Fab-IgG and IgG-Fab expression, HC and LC plasmid were co-transfected using a 1:2 ratio at 1 μg total DNA per 1 mL culture. Transfected Expi293F cells were grown at 37°C in a 5% CO_2_ incubator and shaken at 125 r.p.m. for 3-5 days. Supernatants were harvested by centrifugation at 12,800 g for 10 min, passed through 0.22 μm filters and purified using standard protein A/G chromatography method. Purified Abs were quantified using absorbance at 280 nm along with corresponding protein extinction coefficients.

### Antibody characterization using SDS-PAGE, SEC-MALS and nanoDSF

Purified anti-sperm IgG Abs were first assessed for molecular size by SDS-PAGE. Briefly, 1 μg of protein was diluted in 3.75 μL LDS sample buffer followed by the addition of 11.25 μL nuclease-free water. Proteins were then denatured at 70°C for 10 min in a thermocycler. Next, 0.3 μL of 0.5 M tris (2-carboxyethyl) phosphine (TCEP) was added as a reducing agent to the denatured protein for reduced samples and incubated at room temperature (RT) for 5 min. Bio-Rad Precision Protein Plus Unstained Standard and Novex™ Sharp Pre-stained Protein Standard were used as ladders. After loading the samples, the gel was run for 50 min at a constant voltage of 200 V and washed 3 times with Milli-Q water. Then, the protein bands were visualized by staining with Imperial Protein Stain (Thermo Scientific) for 1 hr followed by overnight de-staining with Milli-Q water. Image J software (Fiji) was used to adjust the brightness and contrasts of the SDS-PAGE gel for visual purposes.

For SEC-MALS experiment, GE Superdex 200 10/300 column connected to an Agilent FPLC system, a Wyatt DAWN HELEOS II multi-angle light-scattering instrument (Wyatt Technology, Santa Barbara, CA), and a Wyatt T-rEX refractometer were used. The flow rate was maintained at 0.5 mL/min. The column was equilibrated with 1X PBS, pH 7.4 containing 200 mg/L of NaN_3_ before sample loading. 50-100 μg of each sample was injected onto the column, and the MALS data were collected and analyzed using Wyatt ASTRA software (Ver. 6).

Next, nanoDSF was used to measure the melting (T_m_) and aggregation (T_agg_) temperatures of the Abs by performing thermal denaturation experiments at the rate of 1°C/min from 25°C to 95°C. The intrinsic tryptophan fluorescence at 330 nm and 350 nm was measured. The melting temperature for each experiment was automatically calculated by Nanotemper PR. Thermcontrol software by plotting the ratiometric measurement of the fluorescent signal against increasing temperature. The aggregation temperature for each experiment was also automatically calculated by Nanotemper PR. Thermcontrol software via the detection of the back-reflection intensity of a light beam that passes the sample.

### Collection and processing of semen samples

Healthy male subjects were asked to refrain from sexual activity for at least 24 hr prior to semen collection. Semen was collected by masturbation into sterile 50 mL sample cups and incubated for a minimum of 15 min post-ejaculation at RT to allow liquefaction. Semen volume was measured, and the density gradient sperm separation procedure (Irvine Scientific) was used to extract motile sperm from liquefied ejaculates. Briefly, 1.5 mL of liquified semen was carefully layered over 1.5 mL of Isolate® (90% density gradient medium, Irvine Scientific) at RT, and centrifuged at 300 g for 20 min. Following centrifugation, the upper layer containing dead cells and seminal plasma was carefully removed without disturbing the motile sperm pellet in the lower layer. The sperm pellet was then washed twice with the sperm washing medium (Irvine Scientific) by centrifugation at 300 g for 10 min. Finally, the purified motile sperm pellet was resuspended in the sperm washing medium, and an aliquot was taken for determination of sperm count and motility using computer-assisted sperm analysis (CASA). All semen samples used in the functional assays exceeded lower reference limits for sperm count (15 × 10^6^ total sperm/mL) and total motility (40%) as indicated by WHO guidelines.[21]

### Sperm count and motility using CASA

The Hamilton-Thorne computer-assisted sperm analyzer (Hamilton Thorne, Beverly, MA), 12.3 version, was used for the sperm count and motility analysis. For each analysis, 4.4 μL of the semen sample was inserted into MicroTool counting chamber slides (Cytonix) and six randomly selected microscopic fields, near the center of the slide, were imaged and analyzed for progressively motile (PM) and non-progressively motile (NPM) sperm count. The parameters that were assessed by CASA for motility analysis were as follows: average pathway velocity (VAP), the straight-line velocity (VSL), the curvilinear velocity (VCL), the lateral head amplitude (ALH), the beat cross-frequency (BCF), the straightness (STR) and the linearity (LIN). PM sperm were defined as having a minimum of 25 μm/s VAP and 80% of STR. The complete parameters of the Hamilton-Thorne Ceros 12.3 software are listed in Table S1.[18,19]

### Whole sperm ELISA

Briefly, half-area polystyrene plates (CLS3690, Corning) were coated with 2 × 10^5^ sperm per well in 50 μL of NaHCO_3_ buffer (pH 9.6). After overnight incubation at 4°C, the plates were centrifuged at the speed of 300 g for 20 min. The supernatant was discarded, and the plates were air-dried for 1 hr at 45°C. The plates were washed once with 1X PBS. 100 μL of 5% milk was incubated at RT for 1 hr to prevent non-specific binding of Abs to the microwells. The serial dilution of mAbs in 1% milk was added to the microwells and incubated overnight at 4°C. Motavizumab, a mAb against the respiratory syncytial virus, was constructed and expressed in the laboratory by accessing the published sequence and used as a negative control for this assay. After primary incubation, the plates were washed three times using 1X PBS. Then, the secondary Ab, goat anti-human IgG F(ab’)_2_ Ab HRP-conjugated (1:10,000 dilutions in 1% milk, 209-1304, Rockland Inc.) was added to the wells and incubated for 1 hr at RT. The washing procedure was repeated and 50 μL of the buffer containing substrate (1-Step Ultra TMB ELISA Substrate, Thermo Scientific) was added to develop the colorimetric reaction for 15 min. The reaction was quenched using 50 μL of 2N H_2_SO_4_, and the absorbance at 450 nm (signal) and 570 nm (background) was measured using SpectraMax M2 Microplate Reader (Molecular Devices, San Jose, CA). Each experiment was done with samples in triplicates and repeated two times as a measure of assay variability.

### Scanning electron microscopy

Briefly, 20 × 10^6^ washed sperm was centrifuged at 300 g for 10 min and the supernatant was discarded without disturbing the sperm pellet. Then, 200 μL of Abs or 1X PBS was added to the sperm pellet, mixed by pipetting and incubated for 5 mins using an end-over-end rotator. Next, 200 μL of 4% PFA was added to the Ab-sperm solution and incubated for 10 min using an end-over-end rotator. 50 μL of fixed sperm samples was filtered and washed through membrane filters (10562, K04CP02500, Osmonics) using 0.15 M Sodium Phosphate buffer. The samples were then dehydrated in a graded series of alcohol, transferred to a plate with the transitional solvent, Hexamethyldisilazane (Electron Microscopy Sciences) and allowed to dry after one exchange. Next, filters were adhered to aluminum stubs with carbon adhesives and samples were sputter-coated with gold-palladium alloy (Au:Pd 60:40 ratio, 91112, Ted Pella Inc.) to a thickness of 3 nm using Cressington Sputter Coater 208 hr. Six random images were acquired for each sample using a Zeiss Supra 25 FESEM (Carl Zeiss Microscopy, White Plains, NY) with an SE2 Electron detector at 2500X magnification.

### Sperm escape assay

Briefly, 40 μL aliquots of purified motile sperm (10 × 10^6^ PM sperm/mL) was transferred to individual 0.2 mL PCR tubes. Sperm count and motility were performed again on each 40 μL aliquot using CASA. This count serves as the original (untreated) concentration of sperm for evaluating the agglutination potencies of respective Ab constructs. Following CASA, 30 μL of purified motile sperm was mixed into 0.2 mL PCR tubes containing 30 μL of Abs or sperm washing medium control. The tubes were then fixed at 45° angles for 5 min at RT. Following this incubation period, 4.4 μL was extracted from the top layer of the mixture with minimal perturbation of the tube and transferred to the CASA instrument to quantify the number of PM sperm. The percentage of the PM sperm that escaped agglutination was computed by dividing the sperm count obtained after treatment with Ab constructs by the original untreated sperm count in each respective tube, correcting for the 2-fold dilution with Ab. Each experimental condition was evaluated in duplicates on each semen specimen, and the average from the two experiments was used in the analysis. Data represents 6 independent experiments with at least n=4 unique semen samples. P values were calculated using a one-way ANOVA with Dunnett’s multiple comparisons test.

### Agglutination kinetics assay

Briefly, 4.4 μL of purified motile sperm (10 × 10^6^ PM sperm/mL) was added to 4.4 μL of Ab constructs in 0.2 mL PCR tubes followed by gentle mixing. Immediately, a timer was started and 4.4 μL of the mixture was transferred to chamber slides. The center field of the slides was then imaged and analyzed by CASA instrument every 30 s up to 90 s. The reduction in the percentage of PM sperm at each time point was computed by normalizing the PM sperm count obtained after Ab treatment to the PM sperm count obtained after control treatment with sperm washing medium. Each experimental condition was evaluated in duplicates on each semen specimen, and the average from the two experiments was used in the analysis. Data represents 6 independent experiments with at least n=4 unique semen samples. P values were calculated using a one-way ANOVA with Dunnett’s multiple comparisons test.

### Fluorescent labeling of sperm

Purified motile sperm were fluorescently labeled using Live/Dead Sperm Viability Kit (Invitrogen, Thermofisher Scientific). SYBR 14 dye, a membrane-permeant nucleic acid stain, stained the live sperm while propidium iodide (PI), a membrane impermeant nucleic acid stain, stained the dead sperm. SYBR14 and PI dye were added to 1 mL of washed sperm resulting in final SYBR 14 and PI concentration of 200 nM and 12 μM respectively and incubated for 10 min at 36 °C. The sperm-dye solution was washed twice using the sperm washing medium to remove unbound fluorophores by centrifuging at 300 g for 10 min. Next, the labeled motile sperm pellet was resuspended in the sperm washing medium, and an aliquot was taken for determination of sperm count and motility using CASA.

### CVM collection and processing

CVM was collected as previously described.[12] Briefly, undiluted CVM secretions, averaging 0.5 g per sample, were obtained from women of reproductive age, ranging from 20 to 44 years old, by using a self-sampling menstrual collection device (Instead Softcup). Participants inserted the device into the vagina for at least 30 s, removed it, and placed it into a 50 mL centrifuge tube for collection. Samples were collected at various times throughout the menstrual cycle, and the cycle phase was estimated based on the last menstrual period date normalized to a 28-day cycle. Samples that were non-uniform in color or consistency were discarded. Donors stated they had not used vaginal products nor participated in unprotected intercourse within 3 days before donating. All samples had pH < 4.5.

### Multiple particle tracking of fluorescently labeled sperm in mucus

Multiple particle tracking was performed as previously described.[12] Briefly, fresh CVM was diluted three-fold using sperm washing medium to mimic the dilution and neutralization of CVM by alkaline seminal fluid in humans and titrated to pH 6.8-7.1 using small volumes of 3 N NaOH. Next, 4 μL of Abs or control (anti-RSV IgG1) was added to 60 μL of diluted and pH-adjusted CVM and mixed well in a CultureWell™chamber slides (Invitrogen, ThermoFisher Scientific) followed by mixing of 4 μL of 1 × 10^6^ fluorescently labeled PM sperm/mL. Chamber slides were incubated for 5 min at RT. Then, translational motions of the fluorescently labeled sperm were recorded using an electron-multiplying charge-coupled-device camera (Evolve 512; Photometrics, Tucson, AZ) mounted on an inverted epifluorescence microscope (AxioObserver D1; Zeiss, Thornwood, NY) equipped with an Alpha Plan-Apo 20/0.4 objective, environmental (temperature and CO2) control chamber, and light-emitting diode (LED) light source (Lumencor Light Engine DAPI/GFP/543/623/690). 15 videos (512 × 512 pixels, 16-bit image depth) were captured for each Ab condition with MetaMorph imaging software (Molecular Devices, Sunnyvale, CA) at a temporal resolution of 66.7 ms and spatial resolution of 50 nm (nominal pixel resolution, 0.78 μm/pixel) for 10 s. Next, the acquired videos were run through a neural network tracking software modified with standard sperm motility parameters to determine the percentage of PM sperm.[23] Data represents 6 independent experiments, each using a unique combination of CVM and semen specimens. P values were calculated using a one-tailed t-test.

### Statistical analysis

All analyses were performed using GraphPad Prism 8 software. For multiple group comparisons, P values were calculated using a one-way ANOVA with Dunnett’s multiple comparisons tests. The comparison between control- and anti-sperm Ab-treated fluorescent PM sperm was performed using a one-tailed t-test. In all analyses, α=0.05 for statistical significance. All data are presented as the mean ± standard error of the mean.

