## Supplementary File for "Engineering Highly Homogenous Tetravalent IgGs with Enhanced Sperm Agglutination Potency"

### Supplementary Information

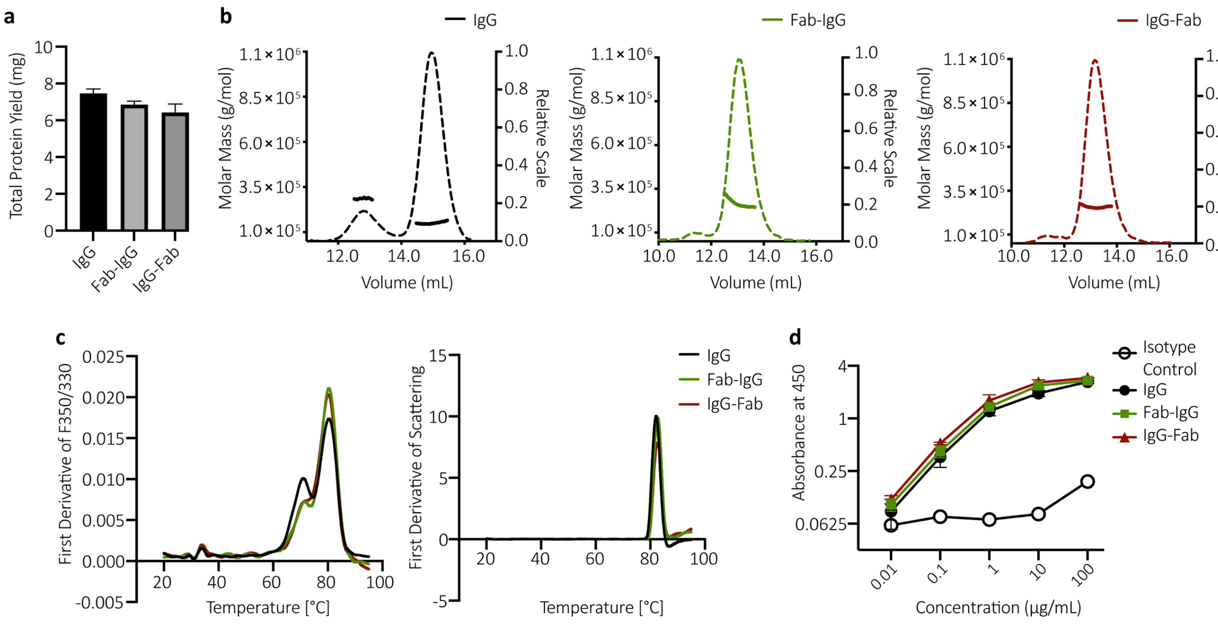


**Figure S1: Additional characterization of tetravalent anti-sperm IgG antibodies.**

(a) Production yield of IgG, Fab-IgG and IgG-Fab purified from 90 mL transfection. Data were obtained from 2 independent transfections. (b) SEC-MALS curves of the indicated Abs; thick lines indicate the calculated molecular mass (left y-axis) and the dotted lines show the homogenous profile (right y-axis) of each antibody. (c) The melting temperatures (left) and aggregating temperature (right) of the indicated Abs as determined by nanoDSF by measuring intrinsic fluorescence and changes in back-reflection of proteins respectively. The experiment was performed in duplicates and averaged. (d) Whole sperm ELISA to assess the binding potency of the indicated Abs to human sperm. Motavizumab (anti-RSV IgG) was used as the isotype control. Data were obtained from n = 3 independent experiments with n = 3 unique semen donors. Each experiment was performed in triplicates and averaged. Lines indicate arithmetic mean values and standard error of mean.

**Table S1. The sperm motility parameters of the Hamilton-Thorne Ceros 12.3.**

| Parameter | Value | Parameter | Value |
|---|---|---|---|
| Frames Per Sec | 60 | Path Velocity (VAP) | 25 µm/s |
| No. of Frames | 60 | Straightness (STR) | 80 % |
| Minimum Cell Size | 3 pixels | VAP Cutoff | 10 µm/s |
| Default Cell Size | 6 pixels | VSL Cutoff | 0 µm/s |
| Minimum Contrast | 80 | Slow Cells | Motile |
| Default Cell Intensity | 20 | Standard Objective | 10X |
| Chamber Depth | 20 µm | Magnification | 1.87 |
